# From scalp to cortex, the whole isn’t greater than the sum of its parts: introducing GetTissueThickness (GTT) to assess age and sex differences in tissue thicknesses

**DOI:** 10.1101/2023.04.18.537177

**Authors:** Sybren Van Hoornweder, Marc Geraerts, Stefanie Verstraelen, Marten Nuyts, Kevin A. Caulfield, Raf Meesen

## Abstract

Noninvasive techniques to record and stimulate the brain rely on passing through the tissues in between the scalp and cortex. Currently, there is no method to obtain detailed information about these scalp-to-cortex distance (SCD) tissues. We introduce GetTissueThickness (GTT), an open-source, automated approach to quantify SCD, and unveil how tissue thicknesses differ across age groups, sexes and brain regions (n = 250). We show that men have larger SCD in lower scalp regions and women have similar-to-larger SCD in regions closer to the vertex, with aging resulting in increased SCD in fronto-central regions. Soft tissue thickness varies by sex and age, with thicker layers and greater age-related decreases in men. Compact and spongy bone thickness also differ across sexes and age groups, with thicker compact bone in women in both age groups and an age-related thickening. Older men generally have the thickest cerebrospinal fluid layer and younger women and men having similar cerebrospinal fluid layers. Aging mostly results in grey matter thinning. Concerning SCD, the whole isn’t greater than the sum of its parts. GTT enables rapid quantification of the SCD tissues. The distinctive sensitivity of noninvasive recording and stimulation modalities to different tissues underscores the relevance of GTT.

## 1. Introduction

Noninvasive techniques to record and modulate brain activity are appealing due to their safety, accessibility, ease-of-use, potential for understanding clinical diagnoses, and therapeutic value. These techniques utilize various mechanisms of action, with some, such as transcranial electrical stimulation (tES) and electroencephalography (EEG), relying on the conduction of electric currents through tissues (Nitsche and Paulus 2000; Nunez and Srinivasan 2006; Bikson et al. 2012; Giordano et al. 2017), and others, such as transcranial magnetic stimulation (TMS) and magnetoencephalography (MEG), using electromagnetic fields (Singh 2014; Klomjai et al. 2015; Fitzgerald and Daskalakis 2022). Still other techniques, such as transcranial focused ultrasound (tFUS) and functional near infrared spectroscopy (fNIRS), rely on the propagation of sound and light waves, respectively (Reich 2005; di Biase et al. 2019).

Irrespective of the mechanism of action, one aspect that all of these techniques share is the need to traverse the tissues in between the modality and the cortex. Consequently, all scalp-based neuroimaging and noninvasive brain stimulation techniques are impacted by variability in the thicknesses of the intermediary tissues. However, there is no easy and fast approach to measure tissue composition. There is a need for a quick and easily implemented method to measure tissue thicknesses, particularly as the effect of tissue thickness variability differs for each modality, depending on the mechanisms of action.

For instance, the total scalp-to-cortex distance (SCD) is important for modalities that rely on magnetic fields, such as TMS and MEG, given that magnetic field strength is largely determined by the distance from the source (Cuffin 1993; Kozel et al. 2000; McConnell et al. 2001; Nahas et al. 2001; Herbsman et al. 2009; Julkunen et al. 2012; Nathou et al. 2015; Hanlon et al. 2019). On the other hand, for modalities that use electric fields, such as tES and EEG, tissue composition is crucial, with bone tissue thickness attenuating the cortical stimulation intensity more than soft tissue thickness due to its higher electrical resistivity (Cuffin 1993; Nunez and Srinivasan 2006; Hagemann et al. 2008; Truong et al. 2013; Opitz et al. 2015; Butler et al. 2019). The volume of grey matter exposed to light emitted by fNIRS is inversely related with SCD, and particularly the thickness of the highly vascularized bone and soft tissue layers strongly affect fNIRS (Rolf and Andrew 2008; Cui et al. 2011; Häußinger et al. 2011; Takahashi et al. 2011; Brigadoi and Cooper 2015). Finally, for tFUS, the thickness of tissues also plays a key role in determining the effects, and the induced sound waves are strongly influenced bone thickness (Guo et al. 2021; Zhang et al. 2021). Thus, it is critical to understand SCD and the tissues comprising SCD to further elucidate the effects of noninvasive brain stimulation and recording.

The goal of this study was to develop and test an automated method of measuring tissue thicknesses in user-defined locations such as EEG coordinates. Prior studies examining SCD and/or the underlying tissues have often resorted to manual measurements (Kozel et al. 2000; McConnell et al. 2001; Stokes et al. 2005; Herbsman et al. 2009; Lu, Chan, et al. 2019; Lu, Lam, et al. 2019; Lu et al. 2021). Besides being labor-intensive, this manual approach provides no information about the tissues comprising SCD. Instead, it only provides a single measure, ignoring the unique properties of the underlying tissue, which are highly relevant for modalities such as tES, EEG, tFUS and fNRIS. In addition, manual measurements typically involve calculating the Euclidean distance between two points on a 2D MRI slice (Kozel et al. 2000; McConnell et al. 2001; Stokes et al. 2005; Herbsman et al. 2009; Lu, Chan, et al. 2019; Lu, Lam, et al. 2019; Lu et al. 2021). Not only does this introduce the risk of human errors, defining a normal vector on a 2D MRI-slice also ignores the reality of the head as a 3D entity.

Thus, there are converging incentives to implement a novel approach, and recent advancements in the computational modeling may provide a solution. Through automated segmentation and meshing of MRI-scans, accurate multi-compartment 3D models of the head are established, consisting of white matter, grey matter, cerebrospinal fluid (CSF), veins, compact bone, spongious bone, muscle, eye and soft tissues (Thielscher et al. 2015; Nielsen et al. 2018; Huang et al. 2019; Puonti et al. 2020). Here, we leverage these advancements to introduce GetTissueThickness (GTT), a vectorized, automated algorithm that informs on the tissues comprising SCD at a user-specified region of interest (ROI), which can be either an MNI or subject coordinate of a grey matter or scalp ROI (Zarei et al. 2013; Provencher et al. 2016).

Using GTT, we investigated sex and age differences in tissue thicknesses across different representative anatomical locations that are often recorded and stimulated using noninvasive techniques. Previous studies have shown variations in all the SCD tissues across women and men, and younger and older adults (Blatter et al. 1995; Akiyama et al. 1997; Kozel et al. 2000; McConnell et al. 2001; Makrantonaki and Zouboulis 2007; Greenberg et al. 2008; Hatipoglu et al. 2008; Sabancıoğulları et al. 2012; Royle et al. 2013; Lillie et al. 2016; Ungar et al. 2018; Anand Meundi and David 2019; Indahlastari et al. 2020; Calisan et al. 2021; McCalley and Hanlon 2021; Hanlon and McCalley 2022). However, these studies typically only investigated the effect of age or sex on a single tissue. We aim to provide a more comprehensive overview of multiple tissue thicknesses simultaneously. Our findings may have important, modality-specific, implications, as different tissue profiles across women and men and younger and older adults may imply systematically different neuroimaging and noninvasive brain stimulation results across these cohorts. We hypothesized that older adults, compared to younger adults, will have decreased soft tissue thickness, increased compact bone, spongy bone and CSF thickness, minor-to-no increases in SCD and a slight decrease in grey matter thickness (Blatter et al. 1995; Akiyama et al. 1997; Kozel et al. 2000; McConnell et al. 2001; Makrantonaki and Zouboulis 2007; Greenberg et al. 2008; Hatipoglu et al. 2008; Sabancıoğulları et al. 2012; Royle et al. 2013; Lillie et al. 2016; Ungar et al. 2018; Indahlastari et al. 2020). Concerning sex, we hypothesized that men would have a generally higher SCD, with smaller between-sex differences in the channels close to the vertex (McCalley and Hanlon 2021; Hanlon and McCalley 2022). We expected men to have thicker soft tissue layers, women to have thicker compact bone layers, and spongy bone, CSF, and grey matter layers to show minor-to-no differences across both sexes (Hatipoglu et al. 2008; Lillie et al. 2016; Anand Meundi and David 2019; Calisan et al. 2021).

## 2. Methods

### 2.1. Participants

We included T1w and T2w structural MRI-scans from younger adults (n = 125, 22–36 years, 72 women) and older adults (n = 125, 65–85 years, 58 women). Data from younger adults were retrieved from the Human Connectome Project dataset (Van Essen et al. 2012). Data from older adults were retrieved from the OASIS3 dataset (LaMontagne et al. 2019). Scan parameters are reported in the original publications of the datasets. MRI-scans were segmented and meshed into finite element head models via the anatomically accurate CHARM procedure (Puonti et al. 2020). The resulting models contained several tissue layers, with the white matter, grey matter, CSF, veins, compact bone, spongy bone, and soft tissue being of particular interest in the context of SCD. Of note, soft tissue not only includes the skin, but also muscles, such as the m. temporalis and m. frontalis, and veins refers to the large veins of the head, such as the superior sagittal sinus. Head mesh quality was visually inspected for accuracy.

### 2.2. GetTissueThickness algorithm

GTT (github.com/SVH35/GetTissueThickness) runs in MATLAB and requires a structure as input, containing several parameters (Table 2). This structure should contain a 3D coordinate, whether this coordinate is situated in the soft tissue (scalp) or grey matter (cortex), the space of the coordinate (subject or MNI space), and whether a visual output should be produced. GTT can be called in MATLAB as follows: GTT(structure). GTT has one dependency: SimNIBS 4, which also contains CHARM, required to create the tetrahedral head models.. Of note, in the following explanation of GTT, a coordinate, fiducial and point are distinguished. A coordinate is a set of three numbers that specify a point in a three-dimensional space. Fiducials and points are special types of coordinates, with a fiducial being a user-defined coordinate (i.e., the coordinate of interest) and a point being a vertex of a tetrahedron / triangle. GTT consists of three steps (Figure 1).

**Figure 1.**
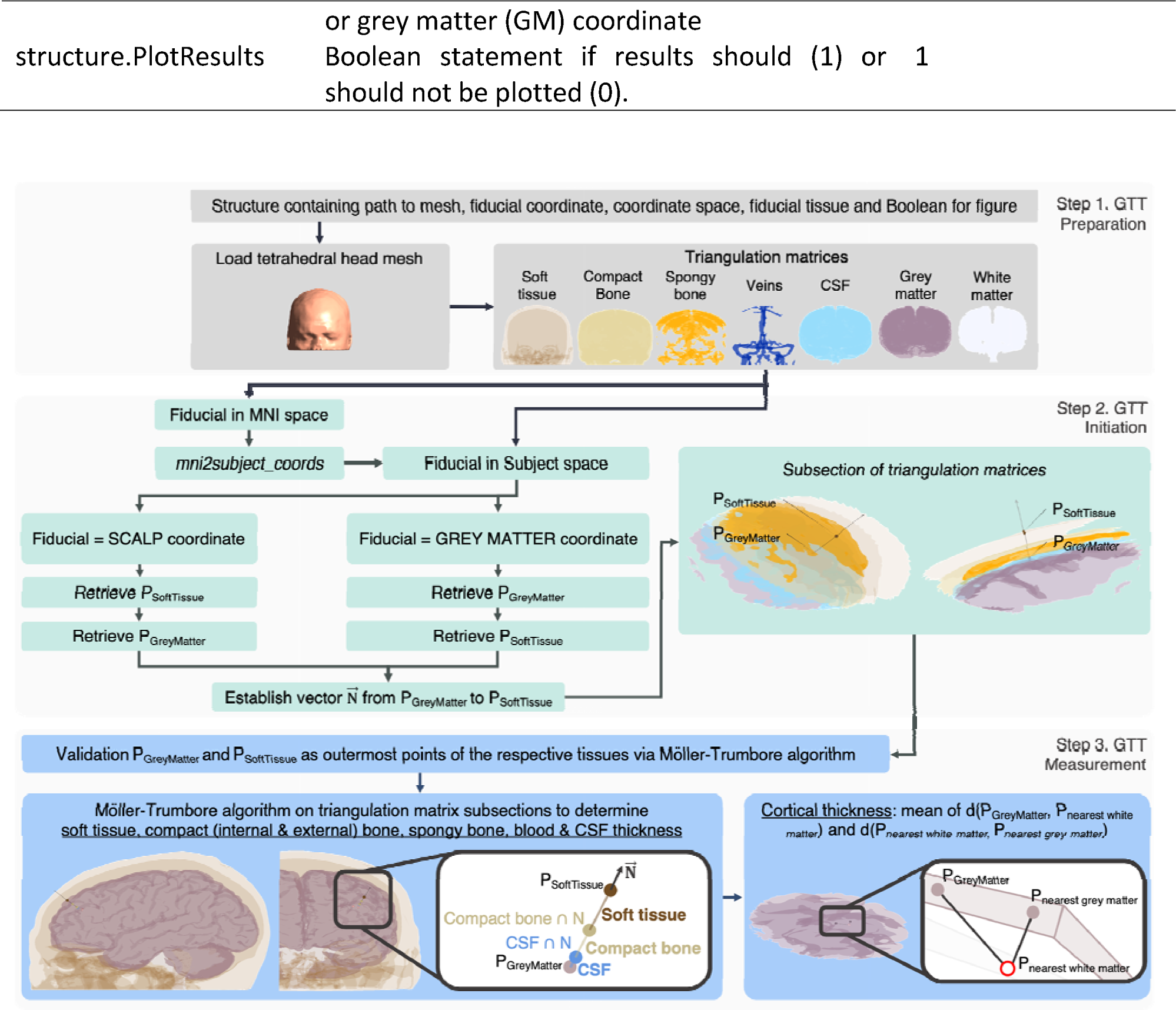
GTT pipeline consisting of three steps: preparation, initiation and measurement.

**Table 2.** GTT input parameters.

| Parameter | Description | Example |
| --- | --- | --- |
| structure.Mesh | Path containing head mesh obtained by CHARM | ... \m2m_subject1 |
| structure.Fiducial | Coordinate of interest | [-52.2 -16.4 57.8] |
| structure.FiducialType | Coordinate space, can be MNI or Subject space | Subject |
| structure.FiducialTissue | Whether structure.Fiducial is a scalp (SCALP) | GM |

#### 2.2.1. Step 1. Preparation

After the head mesh is created via SimNIBS–CHARM, GTT loads the mesh. The mesh is decompiled into triangulation matrices per tissue type (Figure 1), consisting of a point- and connectivity list. The triangulation structure contains the inner and outer layer of each tissue type.

#### 2.2.2. Step 2. Initiation

GTT checks the structure.FiducialType. If set to “MNI”, the fiducial is transformed to Subject space via mni2subject_coords (SimNIBS). GTT then checks structure.BeginTissue. If set to "SCALP", the soft tissue point nearest to the fiducial, P_SoftTissue_, is retrieved and subsequently the grey matter point, P_GreyMatter_, nearest to P_SoftTissue_ is retrieved. If set to "GM", GTT first identifies P_GreyMatter_, and then P_SoftTissue_, as the nearest soft tissue point. Next, a ray, 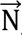, through P_SoftTissue_ and P_GreyMatter_ is established. A sphere is constructed with P_GreyMatter_ as centre, and the r= 2*d(P_GreyMatter_, P_SoftTissue_). This sphere is used to extract a subsection of each tissue’s triangulation matrix, lowering the computational cost of the subsequent procedures.

#### 2.2.2. Step 3. GTT measurement

As aforementioned, the triangulation matrices consist of the inner and outer layer of each tissue. To validate if P_GreyMatter_ and P_SoftTissue_ are both situated on the outer layer, the Möller-Trumbore ray-triangle algorithm is used to assess intersections between a ray and each triangle in a triangulation matrix. The ray is represented by 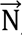, and the triangulation matrix refers to the subsection of the grey matter or soft tissue triangulation matrix. For both the grey matter and soft tissue layers, two intersection coordinates are retrieved. The most outer coordinate is used for the subsequent distance calculations.

Next, the Möller-Trumbore ray-triangle intersection algorithm is performed on the soft tissue, CSF, veins, compact bone and spongy bone layers. Each time, the subsection of the relevant triangulation is tested against 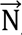, resulting in two coordinates when 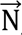 intersects with the tissue: the inner intersection, P_Internal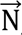_, and the outer intersection, P_External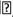_. Via the intersection coordinates, the thicknesses of each tissue are calculated as d(P_Internal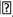_, P_External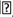_). If a layer of spongy bone is present, the inner and outer compact bone are measured as separate entities. The thicknesses of the CSF, veins, internal compact bone, spongy bone, external compact bone, and soft tissue are summated to obtain the SCD, and compared against the d(P_GreyMatter_, P_SoftTissue_). As both metrics should always correspond, this serves as a validation of the obtained thicknesses.

Considering the convoluted grey matter surface, the grey matter thickness is calculated in line with (Fischl and Dale 2000; Greve and Fischl 2018; Fischer 2022). Specifically, the mean is taken from (1) the shortest distance from P_GreyMatter_ to the white matter, P_nearest white matter_, and (2) the shortest distance from P_nearest white matter_ to the grey matter.

GTT produces a visual output when structure.PlotResults = 1. All used intersection points, 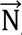, and grey matter, white matter and soft tissue are shown. Besides this, one can also assess the results of GTT by plotting the obtained tissue intersection points on an MRI-scan, as shown in Figure 2. Of note, the accuracy of GTT depends on the accuracy of the supplied mesh. In the case of CHARM, these meshes have been previously validated (Puonti et al. 2020).

**Figure 2.**
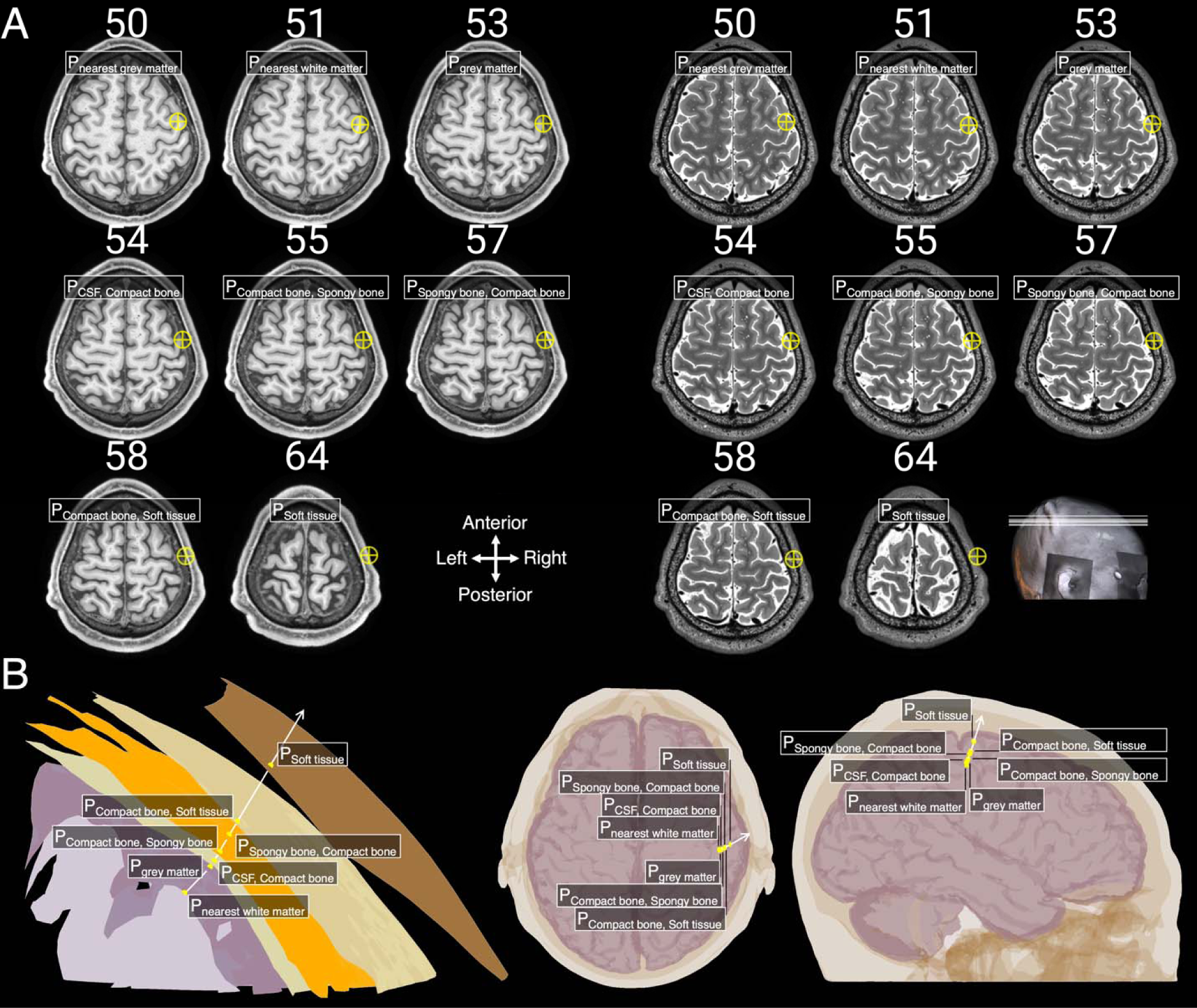
Intersection points of GTT algorithm performed on first younger adult with the coordinates of electrode location C4 supplied. A) intersection points (yellow crosses) plotted on original T1w (left) and T2w (right) MRI-scans of the participant. Numbers above each MRI scan refer to the slice within the MRI scan for this participant. B) Visual output created by GTT.

### 2.3. The effect of aging and sex on head tissues

Previous research has reported differences in the head tissues across younger and older adults, and men and women. However, there has not yet been a comprehensive study of all head tissue thickness changes across ages and sexes. Given the relevance of these tissues for noninvasive brain stimulation and neuroimaging, these differences may induce systemic biases into neuroimaging data and/or noninvasive brain stimulation effects. Therefore, we set out to utilize GTT in 250 healthy adults, and across nineteen EEG 10-20 locations to account for different locations where researchers and clinicians apply noninvasive recordings and stimulation (Okamoto et al. 2004). As the faces of the younger adults in the HCP dataset were blurred to ensure anonymity, soft tissue and SCD were not measured at the blurred regions (i.e., Fp1, Fp2, T3 and T4).

### 2.4. Statistical analyses

Statistical analyses were performed in RStudio (lme4, lm.beta, emmeans) (Bates et al. 2015; Behrendt 2018; RStudio Team 2020; Lenth 2021; R Core Team 2021). For all tests, we used a two-tailed significance value of l1 = 0.05. A linear mixed effects model with random intercepts for PARTICIPANT was constructed per tissue of interest (i.e., SCD, soft tissue, compact bone [i.e., the sum of the internal and external compact bone], spongy bone, and CSF). The fixed effects included SEX (woman / man), AGE GROUP (younger adult / older adult), and CHANNEL (17 to 19 levels). Initially, all interactions, up to the three-way interaction, were included, followed by a stepwise backwards model-building procedure to remove non-significant effects. Interactions present in the final model were analyzed by means of pairwise comparisons, corrected via the Benjamini-Hochberg false-discovery-rate correction for 1,750 to 2,850 pairwise tests, depending on the model.

## 3. Results

### 3.1. GetTissueThickness results

The runtime of GTT depends on several factors (e.g., used computer, if parallel computing is possible in MATLAB and the supplied fiducial). On a MacBook Pro (13-inch, M1, 2020, 8 GB RAM, 8 core CPU) with parallel computing not installed, GTT had a mean runtime of 12.8 ± 5.2 seconds, across ten measurements. Figure 3 provides a global overview of the measured tissues across the different channels, age groups and sexes.

**Figure 3.**
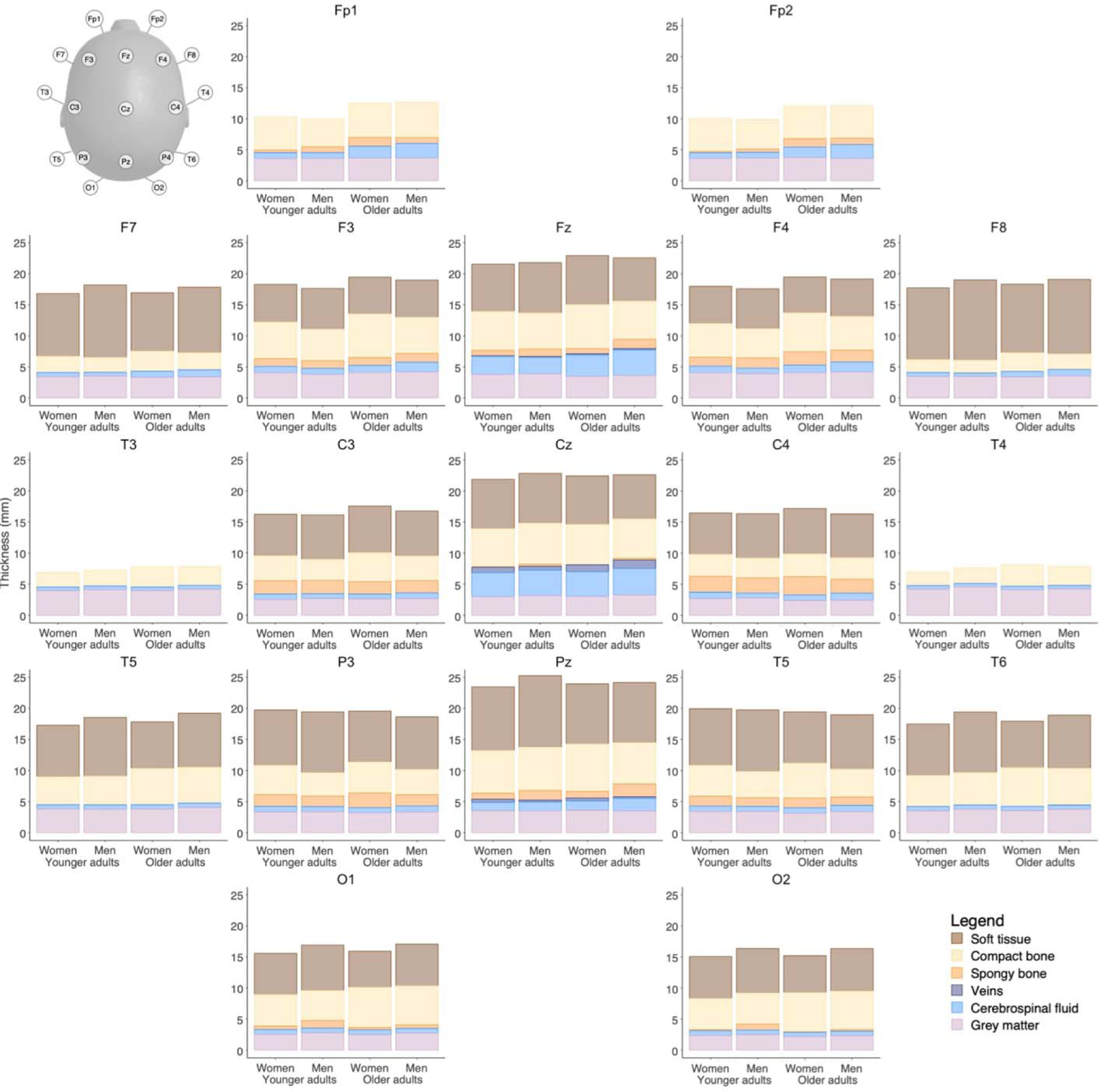
Tissue thicknesses in women and men, and younger and older adults across all brain regions. For Fp1, Fp2, T3 and T4, soft tissue thickness is absent due to the scan anonymization procedure in the HCP dataset used.

### 3.2. Scalp-to-cortex distance

In the SCD model, a significant CHANNEL*SEX (F_14_, _3458_ = 15.170, p < 0.001) and CHANNEL*AGE GROUP (F_14_, _3458_ = 8.83, p < .001) interaction were found. Mean values and differences are shown in Figure 4.

**Figure 4.**
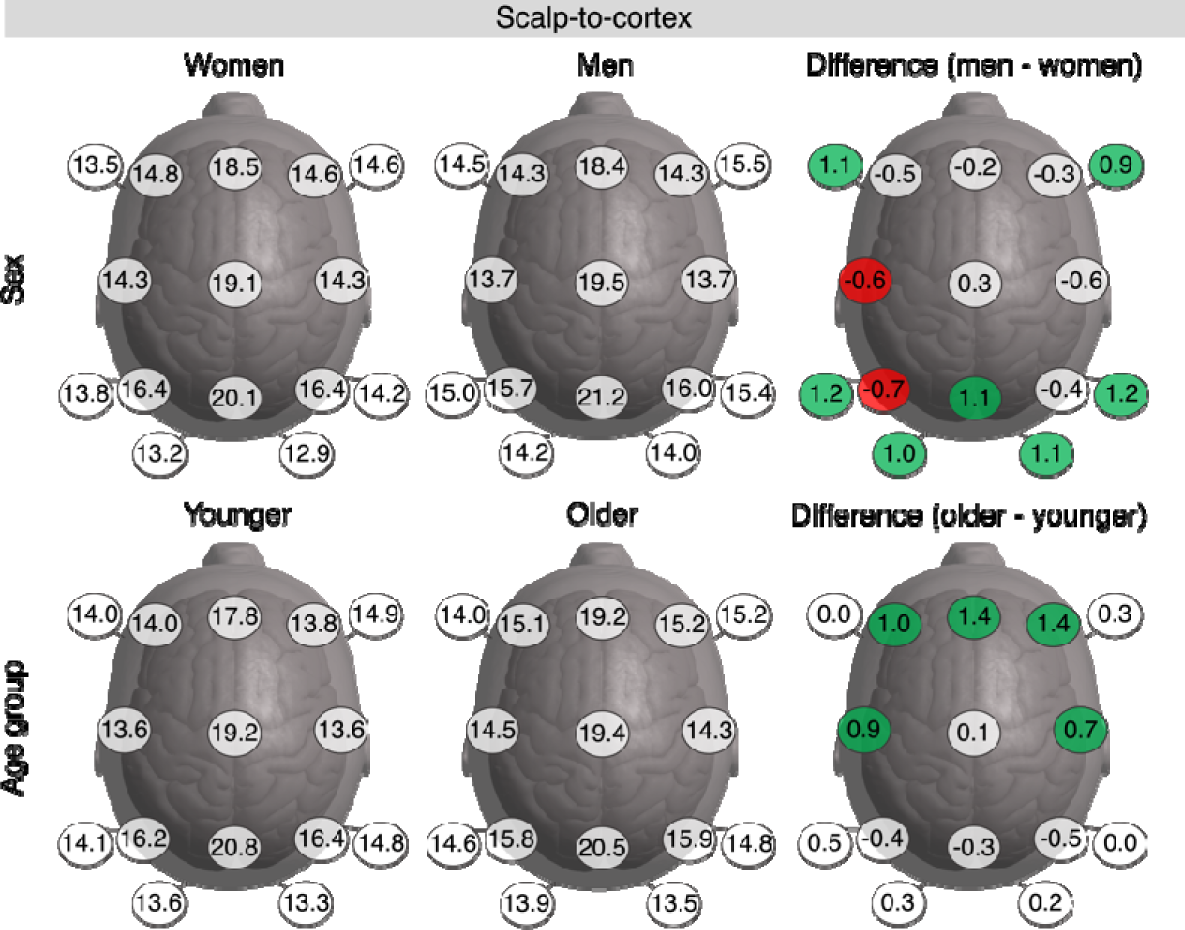
Effect of channel, sex and age group on scalp-to-cortex distance (mm). Colored differences (mm) indicate a significant positive (green) or negative (red) difference.

Concerning the CHANNEL*SEX effect, the effect mostly stems from SCD in the inferior scalp regions being larger in men vs. women, and SCD closer to the midline being more similar across the sexes or even being larger in women. Pairwise comparisons indicated that SCD was significantly larger in men in F7 (z = 3.708, p < 0.001), F8 (z = 3.278, p = 0.002), T5 (z = 4.354, p < 0.001), T6 (z = 4.356, < 0.001), Pz (z = 3.764, p < 0.001), O1 (z = 3.507, p < 0.001), and O2 (z = 3.870, p < 0.001). Conversely, SCD was significantly larger in women in C3 (z = 2.070, p = 0.047) and P3 (z = 2.553, p = 0.014).

Concerning the CHANNEL*AGE GROUP effect, differences in SCD were largest in the frontal and central regions, with other regions showing similar SCD across both age groups. Pairwise comparisons indicated that SCD was significantly higher in older adults in F3 (z = 3.667, p < 0.001), Fz (z = 4.795, p < 0.001), F4 (z = 4.873, p < .001), C3 (z = 3.314, p = 0.001) and C4 (z = 2.539, p = 0.015).

### 3.3. Soft tissue

A significant three-way CHANNEL*AGE GROUP*SEX effect was present (F_14, 3444_ = 3.500, p < 0.001). Mean values and differences are shown in Figure 5. In younger adults, the soft tissue layers were generally thickest in men. Pairwise comparisons indicated that soft tissues were significantly thicker in younger men in F7 (z = 5.331, p > 0.001), F8 (z = 4.690, p > 0.001), T5 (z = 3.984, p = 0.001), P3 (z = 2.876, p = 0.005), Pz (z = 4.288, p < 0.001), P4 (z = 2.694, p = 0.009), T6 (z = 4.884, p < 0.001), and O1 (z = 2.240, p = 0.032), compared to younger women.

**Figure 5.**
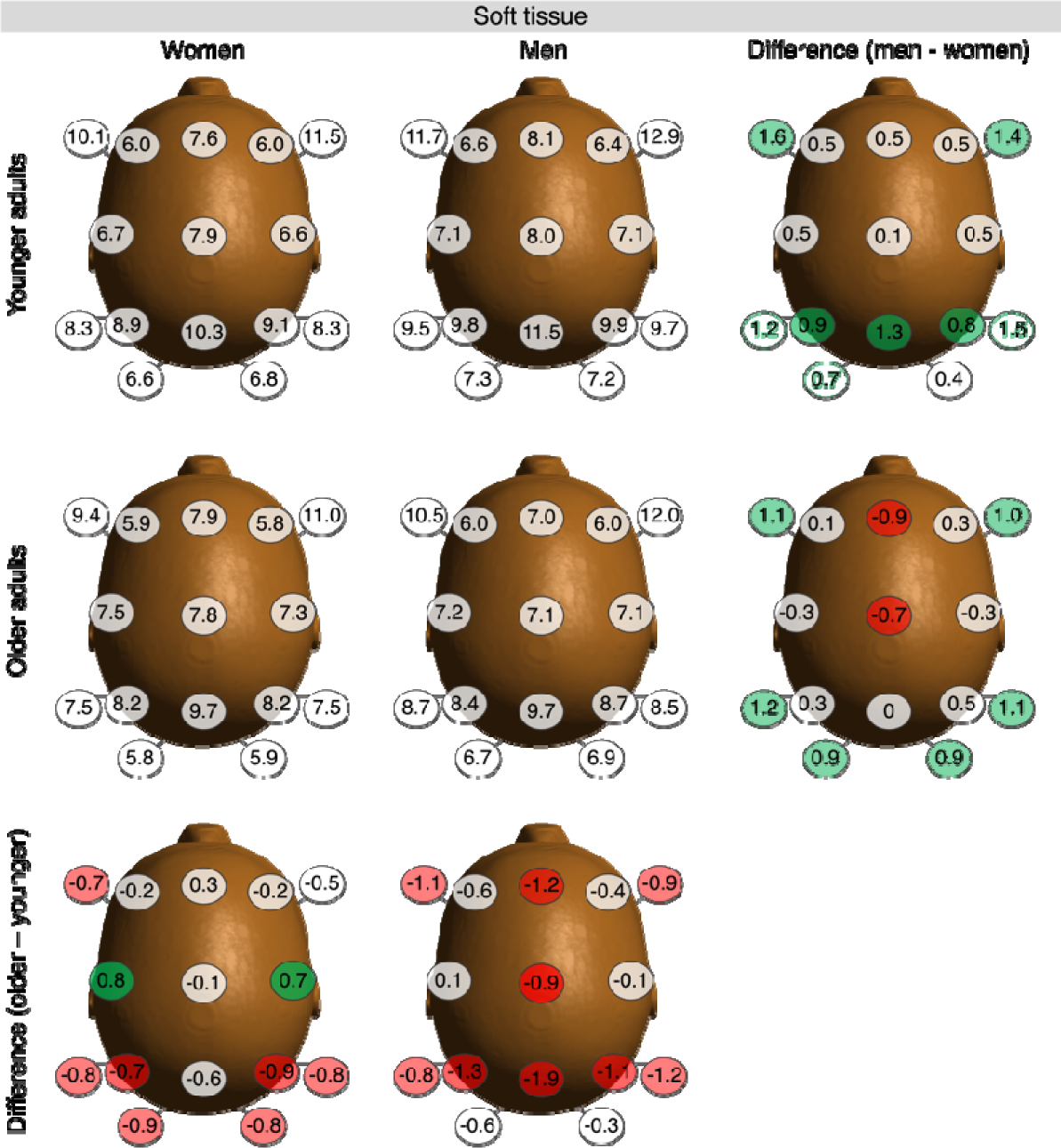
Effect of channel, sex and age group on soft tissue thickness. Colored difference indicate a significant positive (green) or negative (red) difference.

In older adults, the soft tissue layers were also generally thicker in men, though sex differences were less uniform, with the soft tissues in the midline even being thicker in older women. Pairwise comparisons indicated a significantly thicker layer in older men in F7 (z = 3.888, p < 0.001), F8 (z = 3.318, p < 0.001), T5 (z = 4.126, p < 0.001), T6 (z = 3.610, p < 0.001), O1 (z = 3.118, p = 0.003) and O2 (z = 3.152, p = 0.002), compared to older women. Conversely, soft tissues were thicker in older women in Fz (z = −3.133, p = 0.002) and Cz (z = −2.428, p = 0.019), compared to older men.

Compared to younger women, older women mostly showed a decrease in soft tissue thickness, except for the central regions C3 and C4, where tissue thickness was highest in older women. Pairwise comparisons indicated significantly reductions in older women in F7 (z = −2.360, p = 0.023), T5 (z =-2.733, p = 0.008), P3 (z = −2.430, p = 0.019), P4 (z = −2.929, p = 0.005), T6 (z = −2.740, p = 0.008), O1 (z = −2.925, p = 0.005) and O2 (z = −2.819, p = 0.007). In contrast, tissue thickness was significantly increased in older women in C3 (z = 2.818, p = 0.007) and C4 (z = 2.285, p = 0.028), compared to younger women.

Soft tissue thickness decreased more uniformly with age in men. Pairwise comparisons indicated significant decreases in older men in F7 (z = −3.734, p < 0.001), Fz (z = −3.832, p < 0.001), F8 (z = −3.089, p = 0.003), Cz (z = −3.041, p = 0.003), T5 (z = −2.253, p = 0.015), P3 (z = −4.349, p < 0.001), Pz (z = −6.244, p < 0.001), P4 (z = −3.801, p < 0.001) and T6 (z = −3.933, p < 0.001), compared to younger men.

### 3.4. Compact bone

A significant three-way CHANNEL*AGE GROUP*SEX effect was present (F_18, 4428_ = 2.7202, p < 0.001). Mean values and differences are shown in Figure 6.

**Figure 6.**
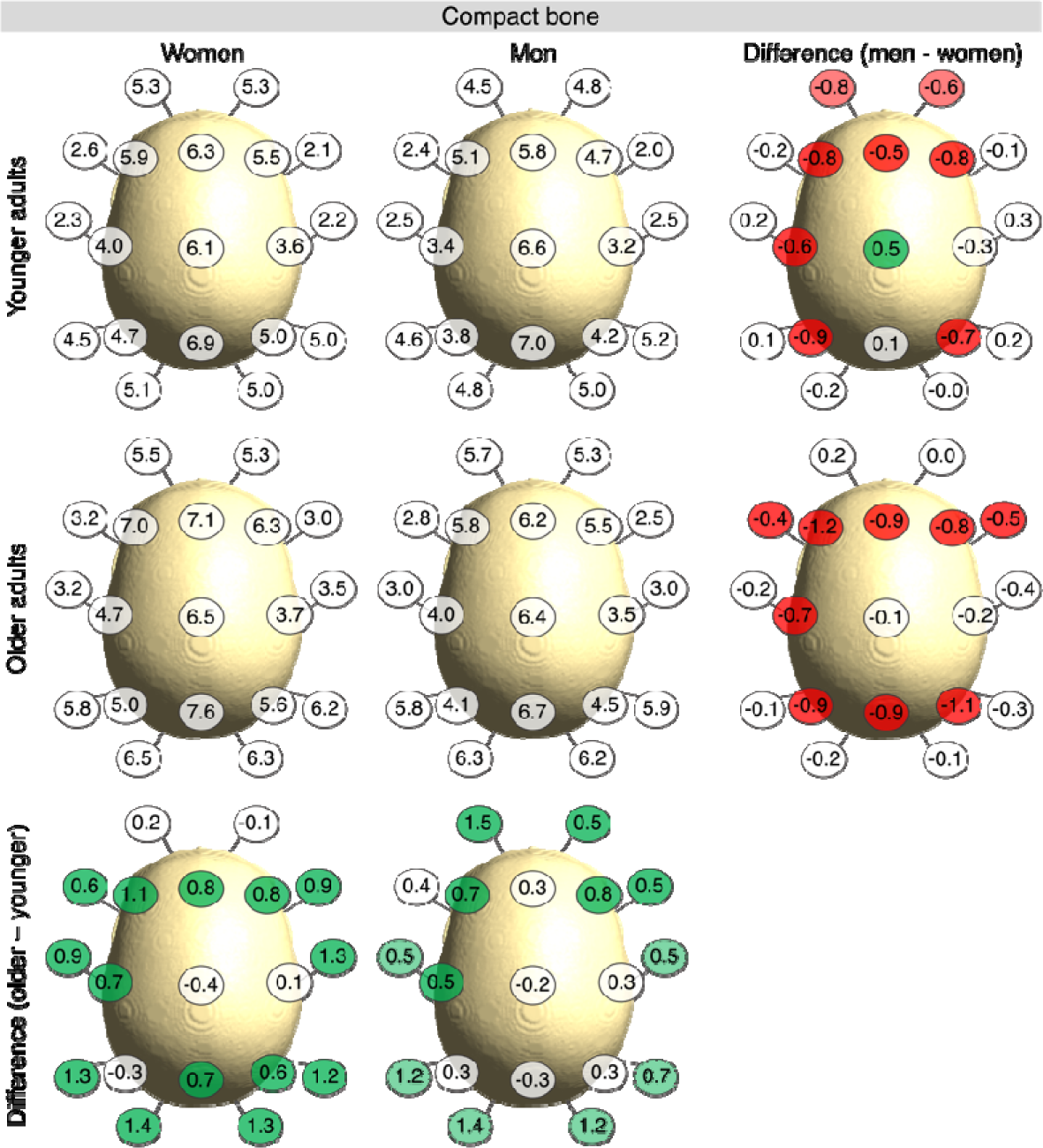
Effect of channel, sex and age group on compact bone thickness. Colored difference indicate a significant positive (green) or negative (red) difference.

In younger adults, the compact bone layer was generally thicker in women. Pairwise comparisons indicated that compact bone tissues were significantly thicker in younger women in Fp1 (z = −3.554, p < 0.001), Fp2 (z = −2.543, p = 0.014), F3 (z = −3.782, p < 0.001), Fz (z = −2.102, p = 0. 043), F4 (z = −3.473, p < 0.001), C3 (z = −2.807, p = 0.006), P3 (z = −4.315, p < 0.001) and P4 (z = −3.406, p < 0.001), compared to younger men. In Cz (z = 2.173, p = 0.036), the bone was thicker in younger men than women.

In older adults, the compact bone was also generally thicker in women, with some regions showing more noticeable sex differences, and other regions showing less noticeable sex differences. Pairwise comparisons indicated that compact bone tissues were significantly thicker in older women in F7 (z = −2.057, p = 0.047), F3 (z = −5.547, p < 0.001), Fz (z = −4.378, p < 0.001), F4 (z = −3.881, p < 0.001), F8 (z = −2.303, p = 0.026), C3 (z = −3.381, p = 0.001), P3 (z = −4.141, p < 0.001), Pz (z = −4.320, p < 0.001), and P4 (z = −5.252, p < 0.001).

Compared to younger women, older women showed an increase in compact bone thickness in most regions. Pairwise comparisons revealed thicker compact bone in older women in F7 (z = 2.958, p = 0.004), F3 (z = 5.204, p < 0.001), Fz (z = 3.901, p < 0.001), F4 (z = 3.904, p < 0.001), F8 (z = 4.385, p < 0.001), T3 (z = 4.264, p < 0.001), C3 (z = 3.081, p = 0.003), T4 (z = 5.886, p < 0.001), T5 (z = 6.223, p < 0.001), Pz (z = 3.502, p < 0.001), P4 (z = 3.036, p = 0.003), T6 (z = 5.704, p < 0.001), O1 (z = 6.668, p < 0.001) and O2 (z = 6.007, p < 0.001), compared to younger women.

Similar to women, compact bone thickness increased with age in men. Pairwise comparisons revealed thicker compact bone thickness in older men in Fp1 (z = 5.227, p = <.0001), Fp2 (z = 2.139, p = 0.0389), F3 (z = 3.331, p = 0.0012), F4 (z = 3.401, p = 0.0009), F8 (z = 2.23, p = 0.0312), T3 (z = 2.348, p = 0.0232), C3 (z = 2.437, p = 0.0183), T4 (z = 2.433, p = 0.0185), T5 (z = 5.254, p = <.0001), T6 (z = 3.184, p = 0.0019), O1 (z = 6.558, p = <.0001) and O2 (z = 5.447, p = <.0001), compared to younger men.

### 3.5. Spongy bone

A significant three-way CHANNEL*AGE GROUP*SEX effect was present (F_18, 4428_ = 2.6269, p < 0.001). Mean values and differences are shown in Figure 7.

**Figure 7.**
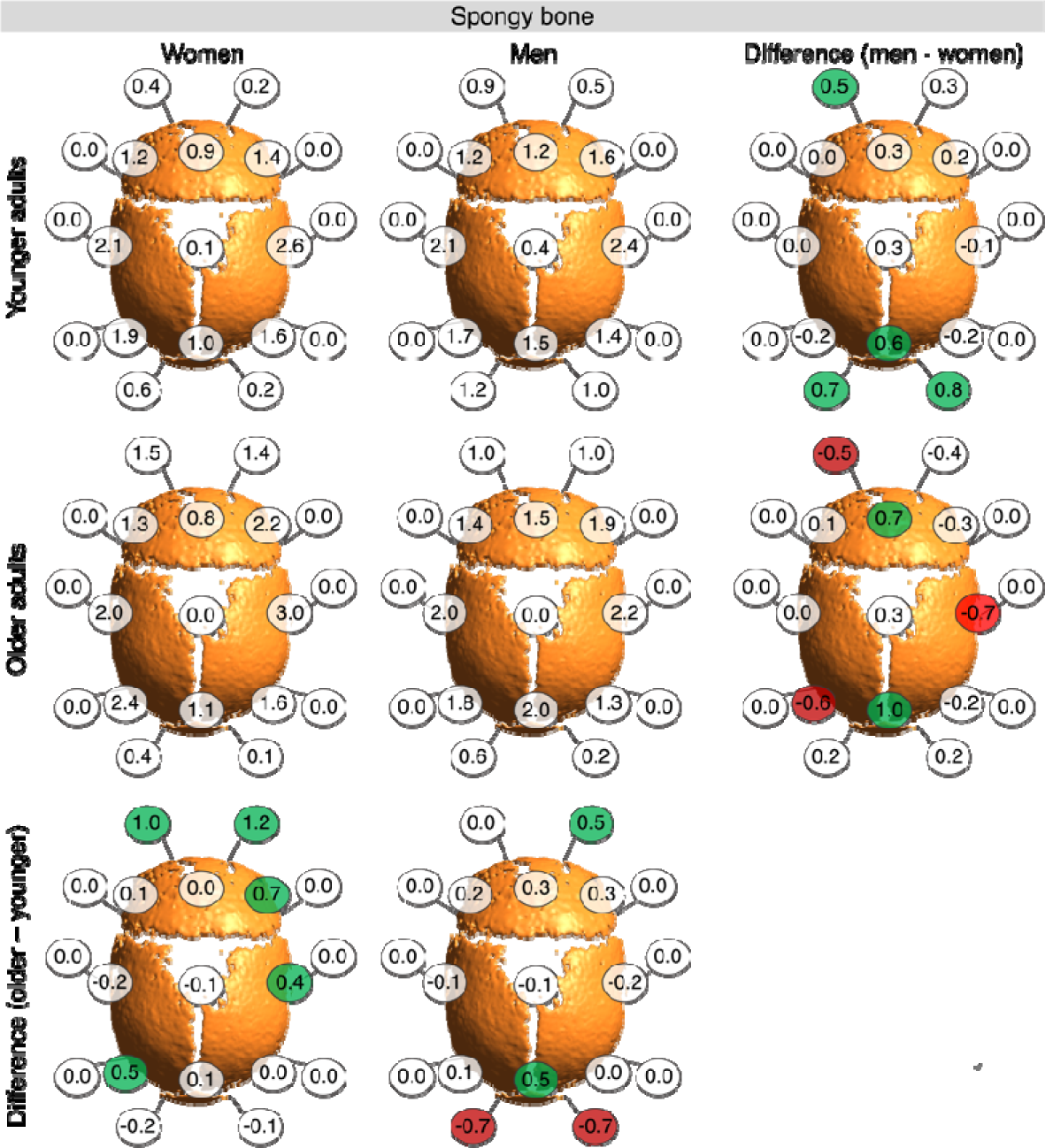
Effect of channel, sex and age group on spongy bone thickness. Colored difference indicate a significant positive (green) or negative (red) difference.

In younger adults, there was a trend towards thicker spongy bone in men compared to women. Pairwise contrasts indicated that spongy bone in Fp1 (z = 2.790, p = 0.008), Pz (z = 3.144, p = 0.002), O1 (z = 3.818, p < 0.001) and O2 (z = 4.477, p < 0.001) were significantly thicker in men, compared to women.

In older adults, the directionality of the effect of sex on spongy bone thickness depended on the location. Pairwise comparisons revealed that older men had larger spongy bone thickness in Fz (z = 3.888, p < 0.001) and Pz (z = 5.505, p < 0.001), compared to older women. Conversely, older women had larger spongy bone thickness in Fp1 (z = −2.840, p = 0.007), C4 (z = −4.130, p < 0.001) and P3 (z = −3.161, p = 0.003), compared to older men.

Compared to younger women, older women generally demonstrated thicker spongy bone layers. Pairwise comparisons revealed significant increases in Fp1 (z = 5.889, p < 0.001), Fp2 (z = 6.683, p < 0.001), F4 (z = 4.232, p < 0.001), C4 (z = 2.155, p = 0.044) and P3 (z = 2.849, p = 0.007).

The effect of age on spongy bone thickness was smaller in men. Pairwise comparisons revealed that spongy bone was thicker in older men in Fp2 (z = 2.883, p = 0.006) and Pz (z = 2.865, p = 0.006), and thicker in younger adults in O1 (z = −3.703, p < 0.001) and O2 (z = −4.100, p < 0.001).

### 3.6. Cerebrospinal fluid

A significant three-way CHANNEL*AGE GROUP*SEX effect was present (F_18, 4428_ = 2.2099, p = 0.002). Mean values and differences are shown in Figure 8.

**Figure 8.**
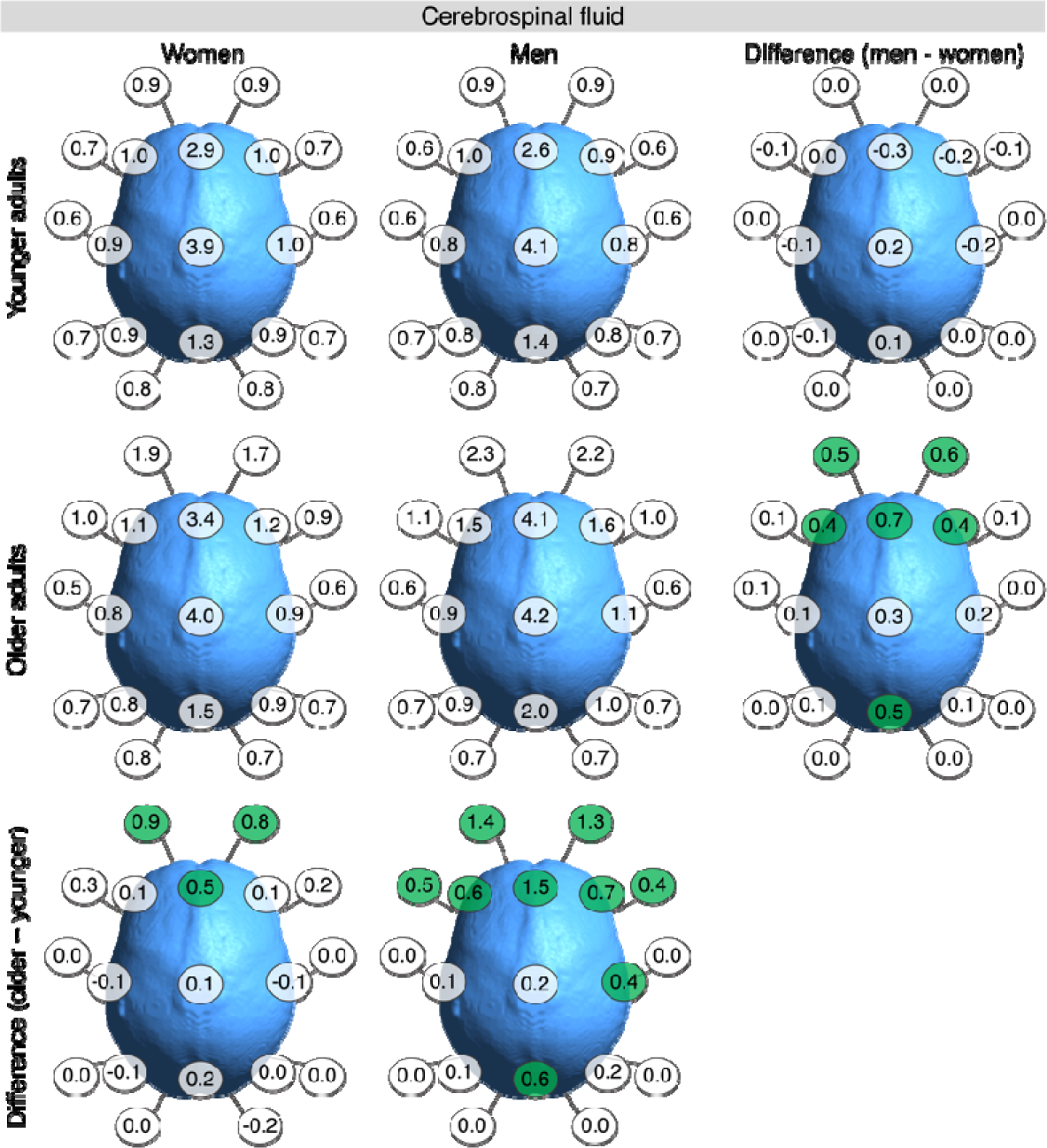
Effect of channel, sex and age group on cerebrospinal fluid thickness. Colored differences indicate a significant positive (green) or negative (red) difference.

In younger adults, there were no significant differences in CSF thickness in women versus men, across all brain regions.

In older men, the CSF was generally thicker than in older women. Post-hoc comparison revealed that this difference was significant in Fp1 (z = 3.606, p < 0.001), Fp2 (z = 4.194, p < 0.001), F3 (z = 3.101, p = 0.004), Fz (z = 5.220, p < 0.001), F4 (z = 3.090, p = 0.005) and Pz (z = 4.036, p < 0.001).

In women, CSF thickness was generally thicker in older individuals. However, age differences were minor overall. Post-hoc comparisons revealed significant increases in older women in Fp1 (z = 7.156, p < 0.001), Fp2 (z = 5.913, p < 0.001) and Fz (z = 3.905, p < 0.001), compared to younger women.

In men, changes across the adult lifespan were more noticeable, with most regions showing increased CSF thickness in older men. Post-hoc comparisons revealed significantly higher CSF thickness in older adults in Fp1 (z = 10.59, p = < 0.001), Fp2 (z = 9.48, p < 0.001), F7 (z = 3.868, p < 0.001), F3 (z = 4.154, p < 0.001), Fz (z = 10.976, p < 0.001), F4 (z = 5.18, p < 0.001), F8 (z = 3.111, p = 0.004), C4 (z = 2.624, p = 0.018), and Pz (z = 4.404, p < 0.001).

### 3.7. Grey matter

A significant CHANNEL*SEX (F_18, 4446_ = 1.6348, p = 0.044) and CHANNEL*AGE GROUP (F_18, 4446_ = 3.4824, p < 0.001) interaction was present. Mean values and differences are shown in Figure 9.

**Figure 9.**
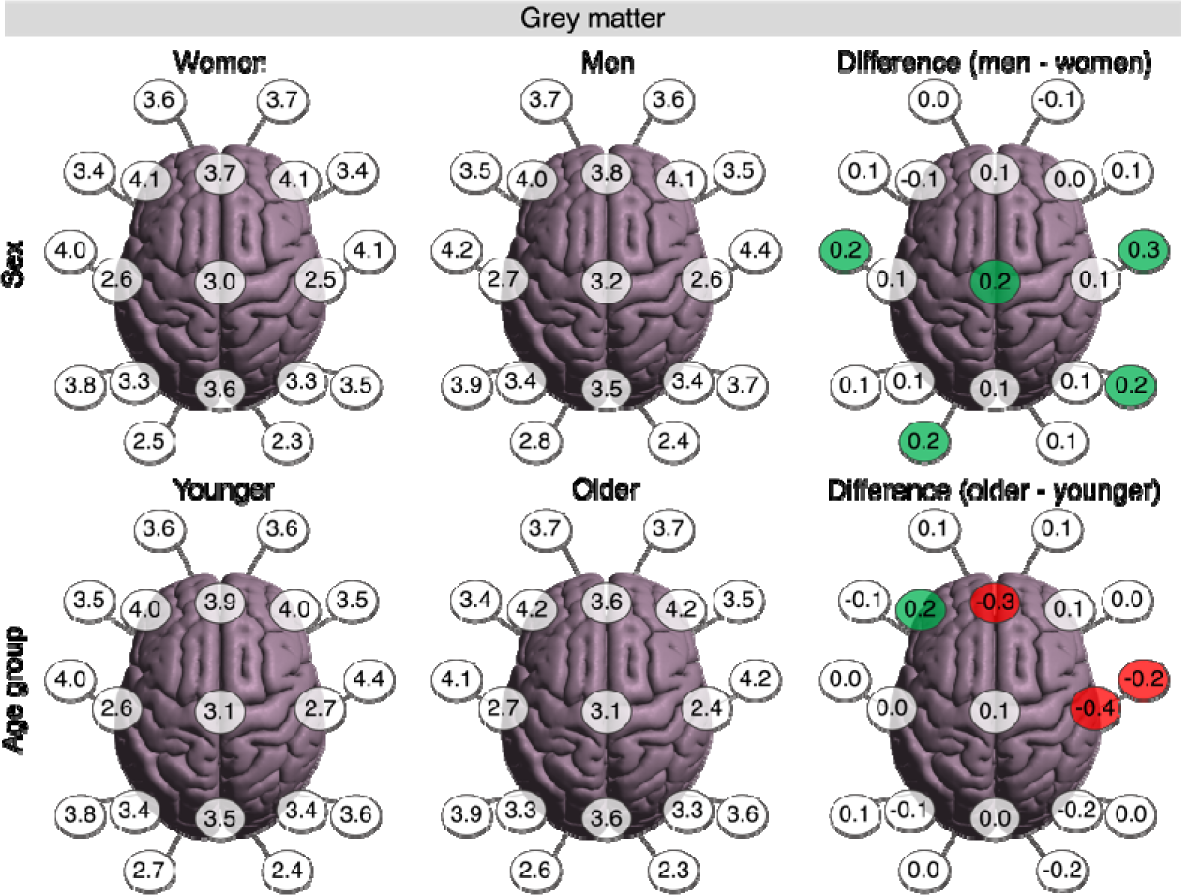
Effect of channel, sex and age group on grey matter thickness. Colored difference indicate a significant positive (green) or negative (red) difference.

Concerning the CHANNEL*SEX interaction, pairwise comparisons indicated that the grey matter was thicker in men, in T3 (z = 2.077, p = 0.047), Cz (z = 2.638, p = 0.011), T4 (z = 2.852, p = 0.006), T6 (z = 2.390, p = 0.022) and O1 (z = 2.680, p = 0.010).

Concerning the CHANNEL*AGE GROUP interaction, a significant decrease in grey matter thickness was present in Fz (z = −3.196, p = 0.002), C4 (z = −4.106, p < 0.001) and T4 (z = −2.442, p = 0.019). Conversely, grey matter was thicker in older adults in F3 (z = 2.492, p = 0.017).

## 4. Discussion

Here, we introduced GTT, a fast, easy-to-use and automated open-source algorithm to measure the tissues making up SCD. Our primary objective with GTT was to design an algorithm surpassing current approaches in terms of comprehensiveness, by not only measuring SCD but also the comprising tissues. Additionally, we have included grey matter measurements in GTT which may help to elucidate age- and diagnosis-related changes in cortical thicknesses. Overall, GTT was fast in determining tissue thicknesses, with an average computation time of only 12.8 ± 5.2 seconds on an average computer (8 GB RAM, 8 core CPU). First, we will discuss the result of GTT related to channel, sex and age group differences. Second, we will discuss implications of our results to the fields of noninvasive brain stimulation and neuroimaging and future proposed applications of GTT.

### 4.1. Tissue thicknesses vary across age groups, sexes and channels

We analyzed tissue thickness variations across nineteen commonly used locations in the neuroimaging and noninvasive brain stimulation fields in 250 healthy adults. Our findings largely support our initial hypotheses. In doing so, they highlight that for SCD, the whole is not greater than the sum of its parts as tissue thicknesses differed across age groups and sexes in ways that partially cancelled each other out upon summation. Thus, SCD alone does not represent reality, in which specific tissue related changes occur across the lifespan, between diagnoses, and between patient demographics (e.g., differing sexes).

Regarding the scalp-to-cortex distance (SCD), we observed significant sex-related SCD differences in regions farther from the vertex, with men having larger SCD values than women. In regions closer to the vertex, no-to-opposite sex-related differences were present. To some extent, our findings agree with recent work from (McCalley and Hanlon 2021), reporting no sex differences in SCD at F3 and C3, and larger SCD in men at Fp1, a region farther away from the vertex. The discrepancies between our results and (McCalley and Hanlon 2021) (e.g., we found that SCD in C3 was larger in women) may stem from differences in samples, as (McCalley and Hanlon 2021) included healthy adults and adults with alcohol use disorders with a mean age of approximately 44 years. Concerning the effect of age, we found that SCD in most channels did not exhibit a significant age effect. However, SCD in F3, Fz, F4, C3, and C4 was greater in older adults, mostly concurring with our initial hypothesis and previous results on SCD and aging, reporting minor-to-no increases (Kozel et al. 2000; McConnell et al. 2001; Lu, Chan, et al. 2019). Soft tissue thicknesses were thinner in older adults. The clearest effect of age group was present in men, who generally showed a thicker soft tissue layer than women. Overall, our results coincide with previous literature reporting an age-related decrease in soft tissue thickness, and thicker soft tissue in men (Makrantonaki and Zouboulis 2007; Ungar et al. 2018; Anand Meundi and David 2019).

Concerning the compact bone, tissue thickness generally increased with age, and was higher in women. Changes in the spongy bone were less straightforward. In women, spongy bone was generally thicker in older adults, whereas the effect of age was less noticeable in men. Concerning sex-differences, men generally had a thicker spongy bone layer when they were young, but in older adults, the sex with the thickest spongy bone layer depended on the channel. The general observation that bone thickness increases with age, and is thicker in women corresponds with (Lillie et al. 2016; Calisan et al. 2021). However, our results differ from Lillie et al. (Lillie et al. 2016) in that we mostly observed compact bone thickness to increase whereas Lillie et al. observed spongy bone thickness increases as the main cause of the age-related increases in bone thickness. This discrepancy likely originates from our models being based on MRI data and the work of Lillie et al. being based on high-resolution computed tomography (CT) scanning, which is better suited to differentiate between the spongy and compact bone tissues.

Concerning CSF, we found an age-related thickening, particularly in men. Previous literature linked this increase in CSF to atrophic processes in brain parenchyma (Blatter et al. 1995; Akiyama et al. 1997; Greenberg et al. 2008; Indahlastari et al. 2020). Sex differences were only present in older adults, with older men having thicker CSF layers. Consistent with previous literature, we observed that differences between both age groups were most noticeable in the prefrontal regions (Anderton 2002; Peters 2006). Under the premise that CSF thickness is an inverse proxy for brain volume, the findings indicating a more pronounced thickening of CSF in men and older men having thicker CSF layers than older women, suggest that age-related brain atrophy is more apparent in men. This is corroborated by several studies reporting more age-related brain atrophy in men, compared to women (Cowell et al. 1994; Coffey et al. 1998; Xu et al. 2000).

Grey matter was thicker in women in a patchwork-like manner. Concerning the effect of age, several channels showed a decrease in grey matter thickness, albeit this decrease was significant in only three channels. Moreover, in F3, grey matter thickness was significantly increased. Recently, (Frangou et al. 2022) investigated cortical thickness in 17,075 persons from four to ninety years old, and demonstrated that the largest decreases in cortical thickness take place during the first two decades, corroborating (Fjell et al. 2015). Subsequently, the rate of grey matter thinning is less apparent throughout the rest of the lifespan. The reason we did not find a widespread age-related thinning of grey matter in our study may be due to two factors: firstly, our exclusion of individuals younger than 22 years old, and secondly, our sample size may not have been large enough to account for the natural variation in grey matter thickness and detect the subtle reduction in grey matter thickness occurring after the first two decades of life. Moreover, GTT is currently designed to derive tissue thicknesses via a single fiducial and ray. In doing so, GTT resembles previous approaches to obtain SCD (Kozel et al. 2000; McConnell et al. 2001; Stokes et al. 2005; Summers and Hanlon 2017; McCalley and Hanlon 2021). Alternatively, one can measure tissue thicknesses across multiple fiducials in close vicinity, which can attenuate variability induced by the convoluted nature of the grey matter cortex. Due to the fast runtime of GTT (∼12 seconds), it is highly feasible to do so with GTT, and this might be viable path for future work interested in grey matter thickness.

### 4.2. Implications of GTT and these changes for noninvasive brain stimulation and neuroimaging

GTT revealed that tissue’s thicknesses differ across sexes, age groups, and channels. Here, we discuss the implications of these findings in light of three neuroimaging and three noninvasive brain stimulation modalities. As all of these modalities operate through distinct mechanisms, they are affected differently by the observed changes in tissue thickness among sexes and age groups. While our discussion is restricted to the impact of tissue thickness, it is worth noting that other parameters, such as tissue conductivity and micro-vascularization, are also expected to vary among age groups, sexes, and channels.

Modalities that rely on an electromagnetic action mechanism, such as TMS and MEG, require careful consideration of SCD as the strength of the magnetic field is largely determined by the distance from the source. Previous TMS research has shown the potential risks of not considering for SCD, as TMS over the dorsolateral prefrontal cortex at an intensity informed by TMS over the primary motor cortex resulted in under-stimulation of the dorsolateral prefrontal cortex due to SCD differences (Nahas et al. 2004; Stokes et al. 2005; George et al. 2010; Caulfield et al. 2021). Moreover, Hanlon and McCalley reported that differential SCD ratios across channels can result in meaningful differences in TMS doses across cohorts (Hanlon and McCalley 2022). We build further on these results, by showing that SCD varies in a channel-specific manner based on age group and sex. Via GTT, future research may inform on how SCD precisely differs across channels, age groups and/or sexes. This information can then be leveraged to control for SCD as a confounding variable in TMS and/or MEG analyses, or to individualize TMS.

For modalities governed by an electric mechanism of action, such as tES and EEG, the tissues comprising SCD, and in particular their electric resistivity, play a more vital role. Here, our observation that even if SCD remains identical, the underlying tissues may be altered is impactful and has implications for previous and future work. For one, Indahlastari et al. (Indahlastari et al. 2020) found an inverse relationship between age-related brain atrophy and electric field strength induced by tES, and conclude that adjusting tES current intensity based on brain atrophy may be necessary for future work. Here, we add on to this work by showing that it is crucial to not only consider brain atrophy but also the tissues that make up SCD when adjusting tES intensity in the context of the aging brain. Electric field modeling software packages like SimNIBS and ROAST can be used to achieve this, and GTT can be a valuable tool to provide insights into how the tissues differ across individuals (Thielscher et al. 2015; Huang et al. 2019). Moreover, we highlight that beyond aging, sex is also a relevant factor to consider. The significance of our findings in relation to EEG research is heavily influenced by the employed EEG analyses. If individualized volume conductor source modeling and/or baseline correction are used, the influence of differences in tissue thickness is reduced. However, in research where either one or both of these factors are absent, as is the case in sensor-level resting-state EEG studies, it is necessary to interpret differences in the recorded EEG signal across age groups and/or sexes with care, as part of the observed differences likely resembles differences in tissue thicknesses and not brain activity. Moreover, even when baseline correction is applied (e.g., (Van Hoornweder, Blanco-Mora, et al. 2022)), differences in tissue thicknesses may be of relevance as they can alter volume conduction effects of the EEG signals registered at the sensor space. As this is outside the current scope, future work should investigate this further. Similarly, it would be insightful if future work investigated how these observed differences in tissue thicknesses precisely impact the electrical fields induced by tES across different brain regions and cohorts.

In fNIRS, previous research revealed that light penetration depth is inversely related to SCD. When SCD is higher, less grey matter volume is exposed to NIR light, mostly due to the highly vascularized bone and soft tissue layers (Rolf and Andrew 2008; Cui et al. 2011; Häußinger et al. 2011; Takahashi et al. 2011; Brigadoi and Cooper 2015). Thus, systemic differences in SCD and the comprising tissues across channels, sexes and age groups are important when interpreting fNIRS. Here also, GTT can enable researchers to control for these factors.

Concerning tFUS, the attenuation of sound pressure caused by SCD tissues is of major concern. It is well known that bone yields a particularly strong effect on tFUS, with recent work showing that bone-induced attenuation can be up-to 4.70–7.06 greater than attention caused by the brain or scalp layer (Guo et al. 2021). Additionally, it has been shown that bone thickness has the strongest effects on the bone-induced attenuation of tFUS (Zhang et al. 2021). In this study, we observed that bone is thinnest in the temporal region (i.e., locations T3 and T4), supporting previous work using the temporal window to place the tFUS transducer to attenuate the detrimental effects of bone (Badran et al. 2020; Guo et al. 2021; na and Wang 2021; He et al. 2022). Furthermore, we observed significant differences in compact bone thickness across many channels, with older women generally having the highest values and younger men having the lowest values. As these differences were also present in the temporal region (i.e., the tFUS application site), the increased thickness of compact bone associated with aging and women is likely to have impact tFUS effects. Future work can further investigate the relevance of these changes on the effects of tFUS by means of GTT.

### 4.3. Limitations

The current study has some limitations. First, despite using T1w and T2w MRI-scans, which improved segmentation accuracy compared to models relying solely on T1w images (Nielsen et al. 2018; Van Hoornweder, Meesen, et al. 2022), investigating the compact and spongy bone layers would be better achieved by means of high-resolution CT. However, high-resolution CT comes at the cost of being more invasive and is generally not available in most neuroscientific studies, which tend to opt for MRI due to its better soft tissue contrasts. In other words, when the full picture is of interest (i.e., all tissues comprising the SCD and the brain itself), GTT and the underlying MRI data may be most desirable. Secondly, although the accuracy of GTT can be checked by plotting the retrieved intersection points on the initial MRI-scans (Figure 2), GTT has not yet been validated directly (e.g., post mortem comparison). Nevertheless, it can be argued that the validity of GTT is contingent on the accuracy of the underlying meshes (SimNIBS – CHARM), which have been validated (Puonti et al. 2020).

## 5. Conclusions

In summary, this study introduced GTT, a computationally fast and automated open-source algorithm that measures and plots the tissues making up SCD in an average of ∼12 seconds on a normal computer. GTT can be used in future work to elucidate neurodegenerative processes, interpret how tissue thicknesses are related to noninvasive brain stimulation and/or scalp-based neuroimaging, and prospectively individualize therapy by accounting for neuroanatomical idiosyncrasies. In addition to introducing the software, we used GTT to examine how age and sex differences affect soft tissue, compact bone, spongy bone, CSF, and grey matter thickness. Our findings indicate that all of these tissues differ in unique ways across locations, both sexes and younger and older adults, which cannot be adequately represented by summating them into SCD. The diverse mechanisms underlying different neuroimaging and noninvasive brain stimulation techniques imply that these modalities are differentially affected by changes in the tissues comprising SCD, underscoring the relevance of GTT. We hope that future research will use GTT to consider tissue thicknesses when conducting and interpreting neuroimaging and/or noninvasive brain stimulation research.

## Data availability

The datasets generated during and/or analyzed during the current study are available from the corresponding author on reasonable request. GTT is available on https://github.com/SVH35/GetTissueThickness.

